# A mathematical approach to correlating objective spectro-temporal features of environmental sounds with their subjective perceptions

**DOI:** 10.1101/085621

**Authors:** Thomas F Burns, Ramesh Rajan

## Abstract

Many studies on the subjective perception of non-linguistic sounds have focused on only a single percept, e.g. pleasantness. In the present study, we have examined three different perception-related factors to also allow us to look at any inter-relationships between them that could be related to objective features. Objective features of the NLSs in this database were calculated and subjective perceptions were recorded from participants. These two elements - objective features and subjective perceptions - were then mapped together using various statistical and mathematical techniques. So as to ground our results in a meaningful context, we chose to map a set of objective features to human percepts which had been used in previous studies of NLS perception, but which had not yet been related back to objective features or combinations thereof.

## 1. Introduction

The objective features of sensory stimuli, in part, contribute to our subjective perceptions, e.g., the chemical structure of an odourant is linked to our perception of its odour (Castro et al. 2013), and the wavelength of light being reflected from an object influences our perception of its colour. Perceiving these differences in the objective features of stimuli enables us to reliably navigate our worlds, e.g. colour perception aids in recognising the difference between foliage and fruit (Osorio & Vorobyev, 1996). However, objective properties do not always map simply to perceptions - e.g., for colour perception, the reflectance of an object and the colours of other objects in a scene are perceptually important, independent of the objective feature (wavelength) (Rüttiger et al. 1999). Complex and incomplete mapping between objective features and subjective perception is also true in other sensory modalities, e.g., auditory pitch perception is not only reliant on sound frequency, but also on timbre (Tramo et al. 2005). To fully unravel such relationships methods like the use of artificial neural networks have also been employed with some success (Shao et al., 2003). Such analytical complexity is despite the fact that these subjective perceptions are otherwise quite simple to understand and report, e.g., “the apple is red, not blue”, “that C# is from a violin, not a piano”. Outside of musical timbre (Grey, 1977; Grey & Grodon, 1978), urgency (Momtahan, 1990; Edworthy et al. 1995; Hellier et al., 1993; Burt et al., 1995; Haas & Edworthy, 1996; Graham, 1999), and identification of materials, e.g. the length of a material being struck and whether it is made of metal or wood (Warren & Verbrugge, 1984; Lakatos et al., 1997), tones (Pollack & Ficks, 1954), or subjects, e.g. the gender of a human walker (Li et al., 1991), little is known about the perceptual mappings of generally complex auditory stimuli and their objective features.

In this study we address this issue for a database of complex sounds that can have important meaning in everyday life. Non-linguistic sounds (NLSs) - e.g., music, a passing bus, snoring - have the advantage of being complex, having meaning and being familiar, but without the confounding semantic and linguistic constraints of language. Attempts have been made to describe how humans identify and remember such non-linguistic (environmental) sounds (e.g., Marcell et al., 2007), albeit not based on objective features of the sounds, and other studies have probed various perceptual properties of NLSs (Halpern et al., 1986; Ballas, 1993; Cycowicz & Friedman, 1998; Kumar et al., 2008; Reddy et al., 2009; Reuter & Oehler, 2011; Singh, 2011; Kirmse et al., 2012; Lewis et al., 2012; Talkington et al., 2012) to make findings such as the importance of spectral features to percepts of unpleasantness in NLSs (Halpern et al., 1986; Cox, 2008; Kumar et al., 2008; Reuter & Oehler, 2011). However, almost all of these studies focused on only a single percept where in the present study we have examined three different perception-related factors, to also allow us to look at any interrelationships between them that could be related to objective features.

Objective features of the NLSs in this database were calculated and subjective perceptions were recorded from participants. These two elements - objective features and subjective perceptions - were then mapped together using various statistical and mathematical techniques. So as to ground our results in a meaningful context, we chose to map a set of objective features to human percepts which had been used in previous studies of NLS perception (Marcell et al., 2007), but which had not yet been related back to objective features or combinations thereof. These percepts were: Complexity, Pleasantness, and Familiarity. In addition, we also noted sound identification accuracy.

In mapping these percepts and the sound identification accuracy to objective features of NLSs, we hypothesised that: (1) the percepts and identification accuracy for NLSs can be quantitatively described by and correlate with objective measurements of specific temporal or spectral contents; and (2) different classes of NLSs would possess unique feature-sets of objective measures which could be related to their perceptual differences or their identification accuracy.

## 2. Methods

### 2.1 NLSs database

Publically available NLSs used in previous research (Marcell et al., 2007) and NLSs from online multimedia archive sources (Shafiro & Gygi, 2004) were combined to create a database of 158 sounds. This database contained 144 distinct sound sources with 14 source exemplars (sounds from an identical source, e.g. snoring, but which are a distinct recordings or events) from a broad range of categories (Appendix I). The first step was to normalize all sounds to the same amplitude so that amplitude differences, affecting audibility and level, did not affect ratings. It is recognized that this procedure equalizes sounds that, in life, may be of unequal level - e.g., the sound of a car revving up would be naturally louder (unless originating from a great distance) and would be, from our experience of environmental sounds, perceived to be louder than the sounds of birds singing. The Complexity of such relationships as a function of our experience with sounds makes it a very difficult factor to control and that may well be the reason why it has not been accounted for in previous studies (e.g., Marcell et al., 2007; Gygi et al. 2007). Here, it was decided that all sounds would be normalized to a standard RMS amplitude before being used for perceptions. We normalized all sounds to the RMS level of the loudest sound in the database, using the CoolEdit 2000 sound program; no other change (e.g., to pitch or rate) was applied.

### 2.2 Psychophysics for rating of subjective perceptions

All psychophysics testing was conducted in a quiet room in an isolated corridor of the Department of Physiology, Monash University. Ethics approval for the collection of this data was obtained from the Monash University Standing Committee on Ethics in Research in Humans.

Twelve normal-hearing observers (7 males, 5 females; mean age 19.8 years, SD 0.57) from the undergraduate student population (all non-musicians) at Monash University, Australia, were tested individually using audiometry to ensure normal hearing thresholds across the range from 500 Hz – 8000 Hz (Rajan & Cainer, 2008). Then, in groups of four, participants listened to the 158 NLSs individually, in groups of 20 sounds at a time with a 1 minute break between groups, so as not to cause fatigue. Using Windows Media Player the sounds were played out as .wav from a Dell Inspiron computer and through an external sound card (Audigy Creative Blaster) to two high-quality speakers, with the four participants organised in a semi-circle facing the speakers, at a distance of about 1 metre from the speakers. After listening to each sound once, the participants were instructed to rate the sound on seven-point Likert scales for ‘Complexity’, ‘Pleasantness’, and ‘Familiarity’ (percepts), and were also asked to name the sound source (Accuracy of Naming) by writing down what they thought the sound was and to rate their confidence in identifying the sounds; no verbal interaction between participants was permitted. Subjects were told: “*Your task is to identify each sound as quickly and accurately as you can. In one or two words please describe what you hear, and write the appropriate response on the blank sheet. Each sheet has blocks of trial numbers from 1 to 20. Sounds will be presented to you in blocks of 20 and after each block you will receive a one minute break. After identifying the sound immediately after it is presented, please rate the sound for Familiarity, Complexity, Pleasantness and confidence on a scale of 1 to 7 (e.g. for Pleasantness 1 = not pleasant at all, 7 = highly pleasant). The sound will only be played once so please listen carefully. The time allocated for each sound is 30 seconds*.” This mimicked the method employed by Marcell et al. (2007). Answers were recorded on sheets of paper marked out in blocks of sounds (Block 1 = first 20 sounds), trial number (trial #1, trial #2, etc.), a blank for the name of the sound and 3 rating scales (Pleasantness, Familiarity, and Complexity).

These subjective ratings and sound identification data were then analysed for correlation with objective measures of the rated NLSs.

### 2.3 Objective measures to define complex waveforms

A wide range of temporal and spectral measures of complex waveforms from various disciplines involving signal processing were considered for inclusion in this study. Particular emphasis was placed on those which measured complexity or have been shown to be related to sound percepts relevant or identical to those used in this study. Measures were selected to include a diversity of possible information (including from different information theoretic viewpoints) without being superfluous or repetitive. Since one percept of interest was ‘Complexity’, we included two entropic measures which measure stimulus ‘complexity’ - sample entropy (Richman & Moorman, 2000; Lake et al., 2002) and permutation entropy (Bandt & Pompe, 2002; Riedl et al., 2013; Zanin et al., 2013), derived from chaos and information theory, respectively. An algorithmic complexity measure was also included, the Lempel-Ziv measure (Ziv J & Lempel A, 1978; Xu et al., 1997; Radhakrishnan & Gangadhar, 1998; Zhang et al., 2000; Wu & Zu, 2001; Zhang et al., 2001; Khalatur et al. 2003; Szczepanski et al. 2003; Watanabe et al. 2003; Huang et al. 2003). The remaining objective measures have been used in previous research on sound identification or perception: peaks-related measures (Gygi et al. 2007); fractal dimension estimates (Spasic et al., 2005; Shibayama, 2006; Raghavendra & Dutt, 2010), using both the Higuchi method (Higuchi, 1988) and the NLD method (Kalauzi et al. 2009); mean spectral centroid (Grey & Grodon, 1978; Shao et al. 2003; Gygi et al. 2007; Maher & Studniarz, 2012); root mean squares (RMSs) of discrete frequency ranges (Halpern et al., 1986; Gygi et al. 2007, Kumar et al., 2008; Reuter & Oehler, 2011); harmonicity (Yumoto et al. 1982; Boersma, 1993; Gygi et al., 2007; Lewis et al., 2012); spectral flatness (Jayant and Noll, 1984; Boersma, 2001); and spectral structure variability (SSV) or index (SSI) (Reddy et al. 2009; Singh, 2011; Talkington et al. 2012).

The HNR measure was calculated for all NLSs using a phonetics research program called Praat (Boersma, 2001) and the remaining 18 selected measures were calculated for all NLSs using MATLAB (MATLAB R2012a, The MathWorks Inc., Natick, MA, United States).

### 2.4 Data analysis

#### 2.4.1 Pair wise regression of the entire dataset of NLSs

To determine whether any of the perception-related factors or any of the objective measures were interrelated, we conducted linear regressions in pair-wise fashion for all combinations of the four perception-related factors: namely, Familiarity, Complexity and Pleasantness, and the Accuracy of Naming. Ratings of Familiarity were highly correlated with ratings of Complexity and with the Accuracy of Naming. We therefore deleted this measure of Familiarity as it did not appear to represent an independent percept, and all further analyses of the three perception-related factors used the percepts of Complexity and Pleasantness, and the recorded Accuracy of Naming. We then examined the pair-wise relationships between each subjective perception-related factor and each of the selected objective measures used to define waveform. Since temporal and spectral domains of sounds appear to be affected differently in different types of hearing loss (Strouse et al., 1998; Probst et al., 2006), and our work may have implications for rehabilitation or testing regimes, we grouped the objective measures into these two categories and conducted the pair-wise analyses between each subjective measure and each objective measure within each category. The resulting regression tables were analysed to make judgements (see Results for criteria) as to which perception-related factors were most independent from the others, and, separately, which objective measures could be chosen as *salient objective measures* based on their independentness from the other objective measures for inclusion in the subsequent, more complex analyses. We conducted linear regressions for all pair-wise combinations between each salient objective measure versus each of the three perception-related factors, namely Complexity, Pleasantness, and the Accuracy of Naming. Significance and correlation coefficients were again calculated for all regressions.

#### 2.4.2 Analyses within categories of sound type

The above-noted analyses on the full dataset of NLSs identified general trends for the relationships between each salient objective measure and each of the three perception-related factors. However, there was still a rather large amount of imperfect mapping between the objective measures and each perception-related factor. To strengthen these analyses, we examined the homogeneity of the NLSs for their objective measures, and for the three perception-related factors, to see if stronger relationships might be found by removing outlier NLSs which did not fit a given trend between an objective measure and each of the three perception-related factors. To objectively and precisely identify these outliers, each NLS was placed into an experimenter-determined sound source category (e.g., Primate sounds, Human non-vocal sounds, Nature sounds, etc. - see Appendix I for full list of categories and segregation of sounds). This allowed us to extract any common objective measures underlying the perception of different types of sounds from within the same sound source category.

We conducted one-way ANOVAs to compare the NLS categories for their differences in the salient objective measures and differences in their perception-related factors. All ANOVAs were tested for heteroskedasticity using Brown-Forsythe tests and pair-wise differences between the categories were found using post-hoc Tukey’s tests. Although the determination of what would be considered ‘homogenous’ would have to be somewhat arbitrary, we counted the number of individual significant differences between the categories across all the ANOVAs to determine if there were more differences than similarities (where a similarity is defined as having no significant difference between two NLS categories). If we found more differences than similarities, removing or separating categories of NLSs from the database might allow us to find more perfect mappings between the objective measures and each perception-related factor. If, on the other hand, more similarities than differences were found, mapping improvements would be unlikely

#### 2.4.3 Multi-variate analyses, Principle Component Analysis, and Agglomerative Hierarchical Clustering for data analyses

In a final analysis, we used complex, multivariate analyses, where *combinations* of salient objective measures were mapped to the three perception-related factors for NLSs. We first conducted multiple linear regressions using combinations of objective measures to map onto the perception-related factors and visualised these relationships with bubble plots and biplots from principal component analysis (PCA). However, as this analysis assumed linear and consistent relationships, we conducted separate analyses which were unconstrained by these assumptions, using agglomerative hierarchical clustering (AHC) with Euclidean distance and Ward’s method, set *a priori*, to divide the database into 10 clusters of NLSs, based on their makeup of objective measures for each domain - spectral and temporal. The 10 clusters were then compared for perception-related factors, using one-way ANOVAs, to determine if dissimilar combinations of objective measures were related to dissimilar perceptual reports. This analysis was repeated for different degrees of clustering, e.g. clustering all NLSs into five clusters, then three clusters, and finally two clusters (based on their similarities and differences in perceptual ratings, Accuracy of Naming, and salient objective measures). In a final analysis, data were reduced in dimensionality via PCA and then the AHC method employed, to corroborate any significant differences found between clusters in the three perception-related factors in the high-dimensional cluster analysis described here.

## 3. Results

### 3.1 Interrelationships between Familiarity, Complexity and Accuracy and between sound Complexity and Accuracy

There were significant inter-relationships between each of three percepts, and the Accuracy of Naming (Table 1; all Pearson correlations p < 0.05). Complexity, Familiarity, and Accuracy of Naming had the most robust inter-relationships (r > 0.7), while Pleasantness had a weaker relationship with the other percepts and a very weak relationship with Accuracy of Naming. Complexity was inversely correlated with Familiarity: the more complex a sound, the less likely it was to be rated as a familiar sound. The Accuracy of Naming was, unsurprisingly, positively correlated with perceptions of Familiarity with the sound, and inversely correlated with perceptions of sound complexity, a novel relationship that has not, to the best of our knowledge, been previously reported.

**Table 1.** Correlation matrix (Pearson) of perceptions and identification accuracy.

|  |  |  |  |  |
| --- | --- | --- | --- | --- |
| Variables | Complexity | Pleasantness | Accuracy | Familiarity |
| Complexity |  | **-0.283** | **-0.701** | **-0.807** |
| Pleasantness | **-0.283** |  | 0.197 | **0.321** |
| Accuracy | **-0.701** | 0.197 |  | **0.735** |
| Familiarity | **-0.807** | **0.321** | **0.735** |  |
| * Note: All Pearson correlations had $p < 0.05$ and **bolded** values indicate highly significant relationships ($\alpha=0.001$). | | | | |

Given that Familiarity was so highly correlated with Accuracy of Naming and Complexity, it provided the least amount of independent variation and so was excluded from subsequent analyses. Although Accuracy of Naming appeared to be dependent on Complexity, it was retained in further analysis, given the novelty and non-intuitive nature of this relationship (i.e., it is unclear why Accuracy of Naming should be inversely correlated with Complexity). Thus, overall, we retained for subsequent analyses three perception-related factors: the two percepts of Complexity and Pleasantness, along with the outcome measure of Accuracy of Naming.

### 3.2 Perceptual relationships with individual objective measures

To characterise the NLSs objectively, we started with 19 objective measures that have been used to characterise complex signals. We calculated these measures for our NLSs and conducted pair-wise linear regression analyses of one measure against another measure, for all combinations of measures (see Supplementary Results #2). As expected, many objective measures which were theoretically related or which measured similar qualities of an NLS were significantly correlated with one another for our NLS database too, e.g. HNR and SFM (r=-0.822); Higuchi and NLD estimates (r=0.812). Hence, to avoid redundancy of information captured by the objective measures and to increase the power of subsequent analyses, we identified a set of *salient objective measures* which (for our dataset) accounted for large proportions of the variance in other objective measures within the same domain (i.e., spectral or temporal) – i.e., we first identified the objective measures that correlated most highly with each other.

Although there are many significant relationships among the measures, some are quite weak (e.g. Higuchi FD estimate and duration, r=0.059) and the choice of measures should include some consideration towards the strength of relationships. Any such consideration would have to be made on an entirely arbitrary basis since the ‘strength’ of a correlation is subjective and relative; we decided that if the correlation between two measures was ≥±0.45, only one of the pair would be included in subsequent analyses. Ten cross-relationships with r≥±0.45 were found among the temporal measures and seven among the spectral measures. Retaining only one from each such cross-relationship reduced the 19 objective measures to a subset of seven *salient objective measures* (four temporal and three spectral; listed in Table 2) which, independent of each other, accounted for a large proportion of the total variance among all of the measures. These seven measures were: the NLDFD, the mean peak relative amplitude, the permutation entropy, and the LZ complexity in the temporal domain and the HNR, the mean spectral centroid, and the 1000-2000 Hz RMS contents’ relative amplitude, in the spectral domain.

**Table 2.** Correlation matrix (Pearson) for Complexity, Pleasantness, and Accuracy of Naming, showing their individual, pair-wise relationships with each of the salient measures.

| Salient measures | Accuracy | Complexity | Pleasantness |
| --- | --- | --- | --- |
| NLDFD | **-0.254** | **0.265** | -0.057 |
| Mean peak | -0.038 | -0.039 | **0.324** |
| Permutation entropy | -0.031 | 0.106 | -0.034 |
| LZ complexity (differential binary) | -0.11 | **0.162** | -0.048 |
| HNR | **0.182** | **-0.248** | 0.109 |
| Mean spectral centroid | **-0.35** | **0.293** | -0.089 |
| 1000-2000Hz RMS | 0.076 | -0.052 | -0.021 |

\***Bolded** values indicate significant relationships (at $\alpha=0.05$ )

These *salient objective measures* were subsequently used to correlate with the two retained percepts (Complexity and Pleasantness) and with the Accuracy of Naming of the NLSs. The outcome of these correlations is shown in Table 2. The mean spectral centroid was the best individual descriptor for both Accuracy of Naming (p<0.0001) and Complexity (p=0.0002). The only salient measure to individually correlate significantly with Pleasantness was the temporal measure of mean peak relative amplitude (p<0.0001), which also did not significantly correlate with the Accuracy of Naming or Complexity.

Two measures - one temporal (permutation entropy) and one spectral (1000-2000Hz RMS) - did not correlate significantly with any of Complexity, Pleasantness, or Accuracy of Naming. Note also that whenever a measure was significantly correlated with Accuracy of Naming, it was also significantly correlated with Complexity, except in the case of the temporal measure of LZ complexity (differential binary), which was only significantly correlated with Complexity (p=0.0413).

In summary, there was no strong correlation between any single salient objective measure and the perceptions of Complexity and Pleasantness of NLSs or the Accuracy of Naming the NLSs.

### 3.3 Analysis of NLSs within categories: reducing database homogeneity

The absence of strong correlations between any single salient objective measure and the perceptions and the Accuracy of Naming could be due to heterogeneity in the NLSs database. The NLSs span a wide variety ranging from machine noises to primate and animal vocalisations, and significant heterogeneity in the overall database may warrant analysing NLSs in smaller, alike groups.

For categorisation of the NLSs we initially considered using a naïve group of subjects to listen to the NLSs and generate their own categories (c.f., Marcell et al., 2007). However, this method produces no less heterogeneity since (Marcell et al., 2007) subjects use widely varying methods, ranging from acoustic similarity to sound imagery or sound source for self-categorization of NLSs, producing widely varying categories even for the same set of NLSs; further, even under the same experimental conditions they can produce many nonsense or obscure categories. Hence we adopted a single categorisation rule, using the source of the NLS as the sole basis for categorisation, and implemented by the two experienced experimenters independently and then consultatively if there were any disagreements. The categories selected were: primate (n=14 sounds), non-primate animal (n=38 sounds), tool/machine (n=28), non-animal nature (n=11), human non-vocal (n=6), music (n=21), insect (n=5), explosions/guns (n=10), and uncategorised or other (n=25; this reflected that some NLSs did not fit well into any of the other categories). The allocation of our NLSs to these categories is detailed in Appendix I.

One-way ANOVAs with post hoc Tukey’s tests revealed significant (p<0.05) differences between the NLS categories for percepts and salient objective measures (Appendix III). Perceptual differences between the NLS categories were found for Accuracy of Naming, Complexity, and Pleasantness. For the salient objective measures, differences between the NLS categories were found for permutation entropy, LZ complexity (differential binary), and HNR, but not for NLD fractal dimension, mean peak amplitude, 1000-2000Hz RMS, or mean spectral centroid. The most prominent were for Pleasantness and HNR.

These differences represented only ¼ of the total possible variation or number of potential pair-wise differentiation among the NLS categories for individual percepts or objective measures, i.e. the NLS categories had fewer significant differences than there were potential for (by a factor of three) in perceptual ratings, Accuracy of Naming, and the salient objective measures. Further, the differences between NLS categories for perceptual ratings and Accuracy of Naming were inconsistent between the same NLS categories for salient objective measures.

These analyses do not provide any support that the NLSs database should be treated as categories of sounds, based on sound source. Hence it will be assumed that the NLSs can be treated as a single group of sounds which have an acceptable homogeneity. It is recognised that there were some perceptual and objective differences between the categories vis-à-vis Pleasantness and HNR and for this reason, in relevant subsequent analyses, information on the source of the sound (the category) was retained as supplementary information to assist with describing any trends by sources.

### 3.4 Multivariate analysis

Given that pair-wise regressions of even the reduced subset of most salient objective measures, against any of the three perception-related factors, did not yield strong relationships, we examined if combinations of objective measures would yield better predictions.

#### 3.4.1 Pairs of measures

Multiple linear regression using *pairs of measures* to explain each of the percepts of Complexity and Pleasantness and the Accuracy of Naming the NLSs, provided more significant relationships (c.f., Table 3 vs Table 2). In keeping with the distinction made previously between temporal and spectral domains, we only combined salient measures from the same domain (spectral or temporal). These analyses are summarised in Table 3 and plotted as bubble plots in Figure 1 for the regressions with the highest explanatory power for each percept and Accuracy of Naming.

**Table 3.**
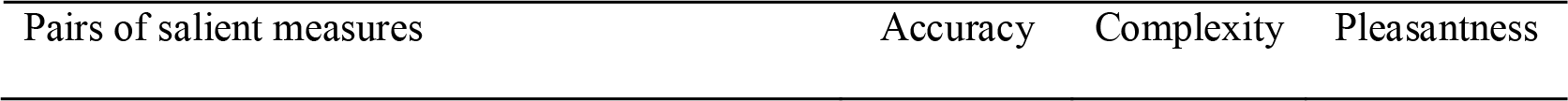
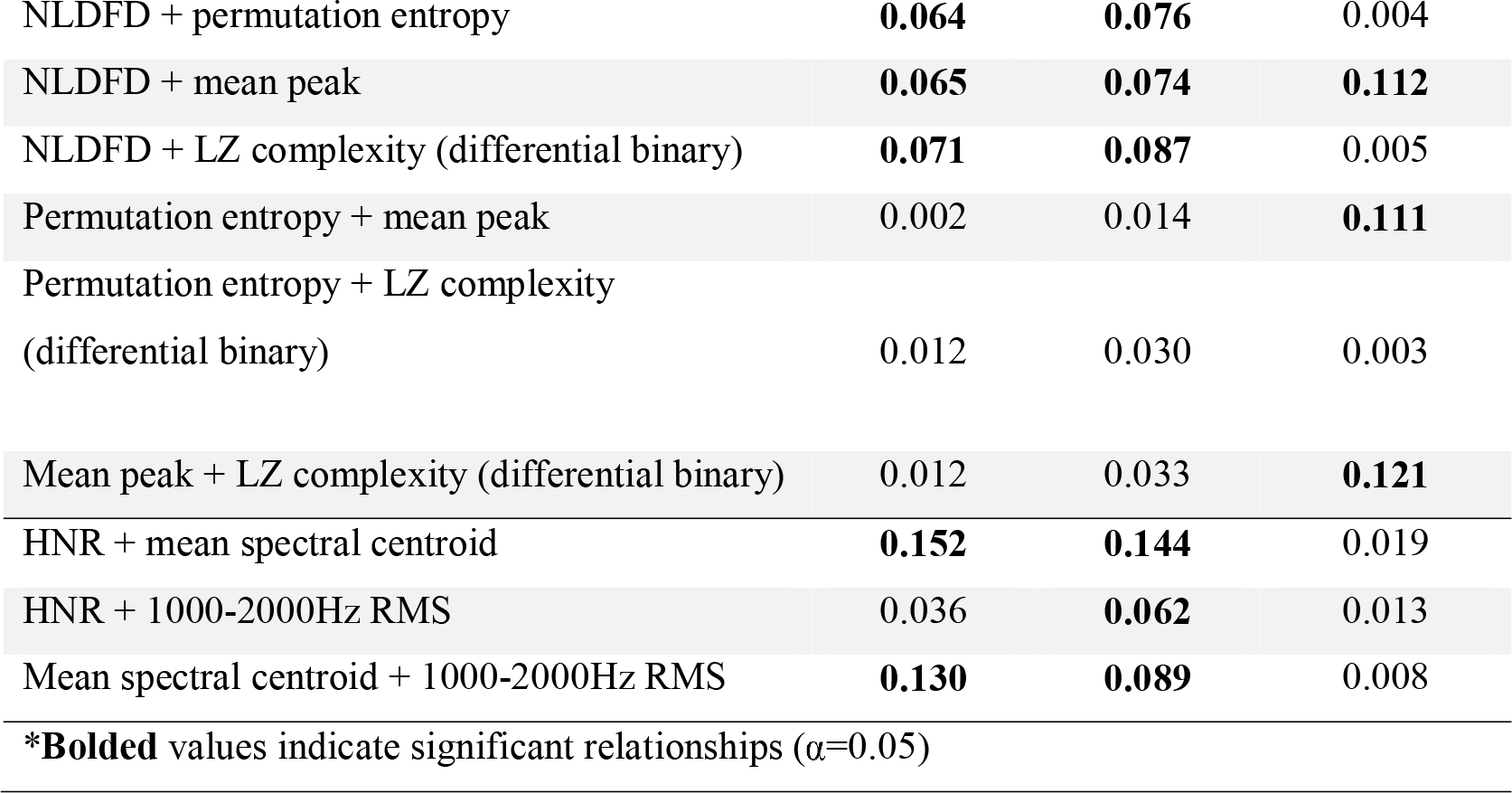
Correlation matrix (R^2^) for Complexity, Pleasantness, and Accuracy of Naming, showing relationships with pairs of salient measures.

| Pairs of salient measures | Accuracy | Complexity | Pleasantness |
| --- | --- | --- | --- |
| NLDFD + permutation entropy | **0.064** | **0.076** | 0.004 |
| NLDFD + mean peak | **0.065** | **0.074** | **0.112** |
| NLDFD + LZ complexity (differential binary) | **0.071** | **0.087** | 0.005 |
| Permutation entropy + mean peak | 0.002 | 0.014 | **0.111** |
| Permutation entropy + LZ complexity<br>(differential binary) | 0.012 | 0.030 | 0.003 |
| Mean peak + LZ complexity (differential binary) | 0.012 | 0.033 | **0.121** |
| HNR + mean spectral centroid | **0.152** | **0.144** | 0.019 |
| HNR + 1000-2000Hz RMS | 0.036 | **0.062** | 0.013 |
| Mean spectral centroid + 1000-2000Hz RMS | **0.130** | **0.089** | 0.008 |
| * **Bolded** values indicate significant relationships ( $\alpha=0.05$ ) | | | |

**Figure 1.**
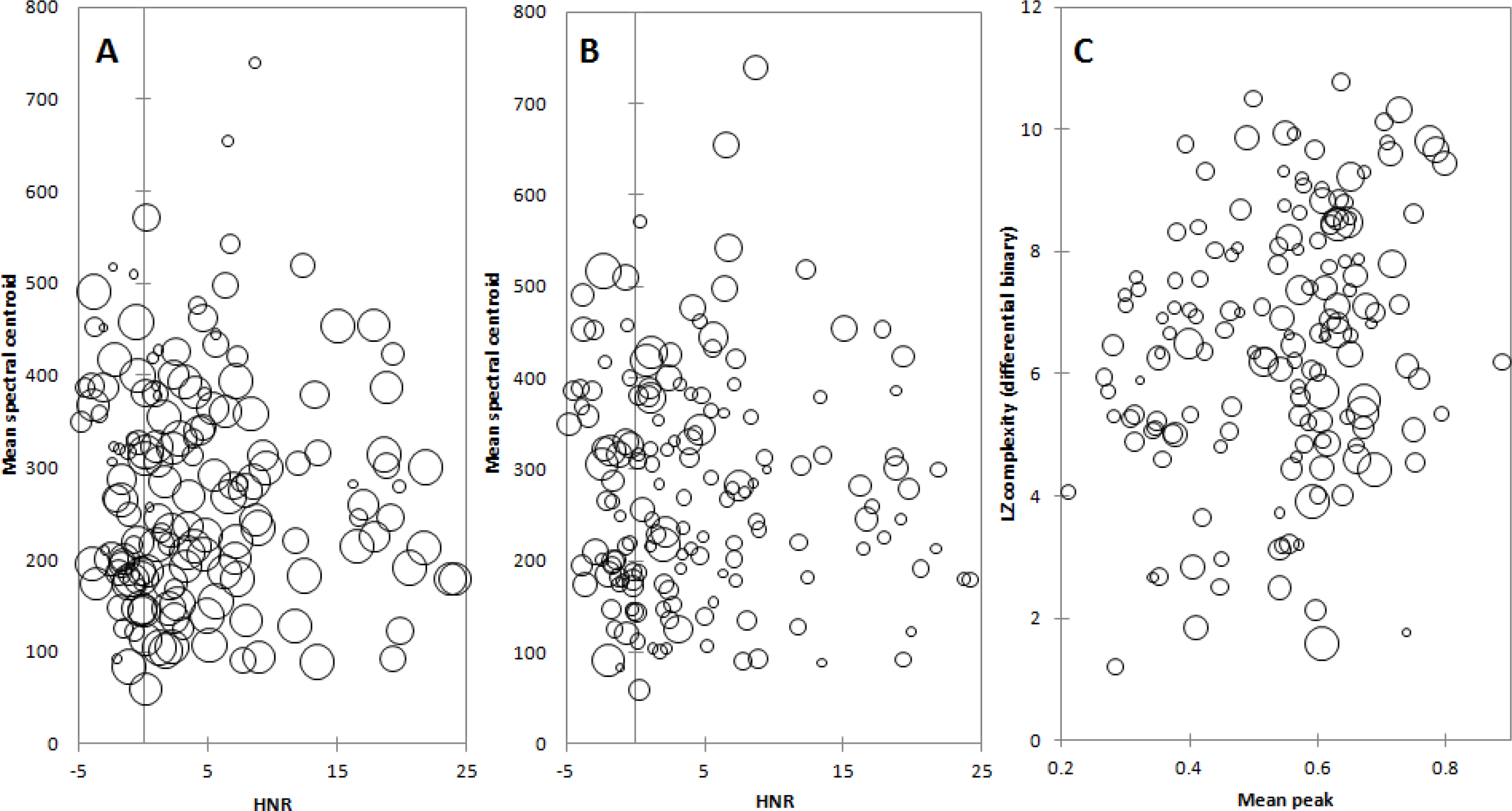
**A:** Mean spectral centrcid and HNR measures bubble plot, where the size of a bubble (representing an Individual NLS) Is scaled to its <u>accuracy of naming</u>. **B:** Mean spectral centroid and HNR measures bubble plot, where the size of a bubble (representing an individual NLS) is scaled to its <u>complexity</u>. C: Mean peak and LZ complexity (differential binary) measures bubble plot, where the size of a bubble (representing an individual NLS) is scaled to its <u>pleasantness</u>.

The spectral combination of HNR and mean spectral centroid provided the largest explanation of both Accuracy of Naming (Figure 1A; R^2^=0.152; p<0.0001) and Complexity (Figure 1B; R^2^=0.144; p<0.0001) of the pair-wise combinations. NLSs which were less frequently identified correctly also tended to have a lower HNR value (Fig. 1A); equally, NLSs with lower HNRs were more frequently rated as having high Complexity (Fig. 1B). Thus, low HNRs appear to be a consistent factor accounting for the above-noted inverse inter-relationship between Complexity and Accuracy of Naming, the latter being a novel relationship that not previously reported.

Mean spectral centroid also appeared to separate NLSs broadly - more complex and less well identified NLSs tended to have higher mean spectral centroids (Fig. 1B).

The temporal combination of mean peak relative amplitude and LZ complexity (differential binary) had the most explanatory power for the perception of Pleasantness of NLSs (Figure 1C; R^2^=0.121; p<0.0001). NLSs with a high Pleasantness where situated mainly on the higher end of the mean peak relative amplitude axis. However, there was no clear distinction for bubble size along the LZ complexity (differential binary) axis, where there seemed to be a fairly equal representation of pleasant and unpleasant NLSs.

#### 3.4.2 Three or more measures

Adding additional objective measures into the multiple linear regressions marginally increased the predictive power for each domain of the three perception-related factors (Table 4 c.f. Table 3), suggesting that, when assuming linearity in the analysis, we had reached the limit of our predictive capabilities using just the salient measures.

**Table 4.**
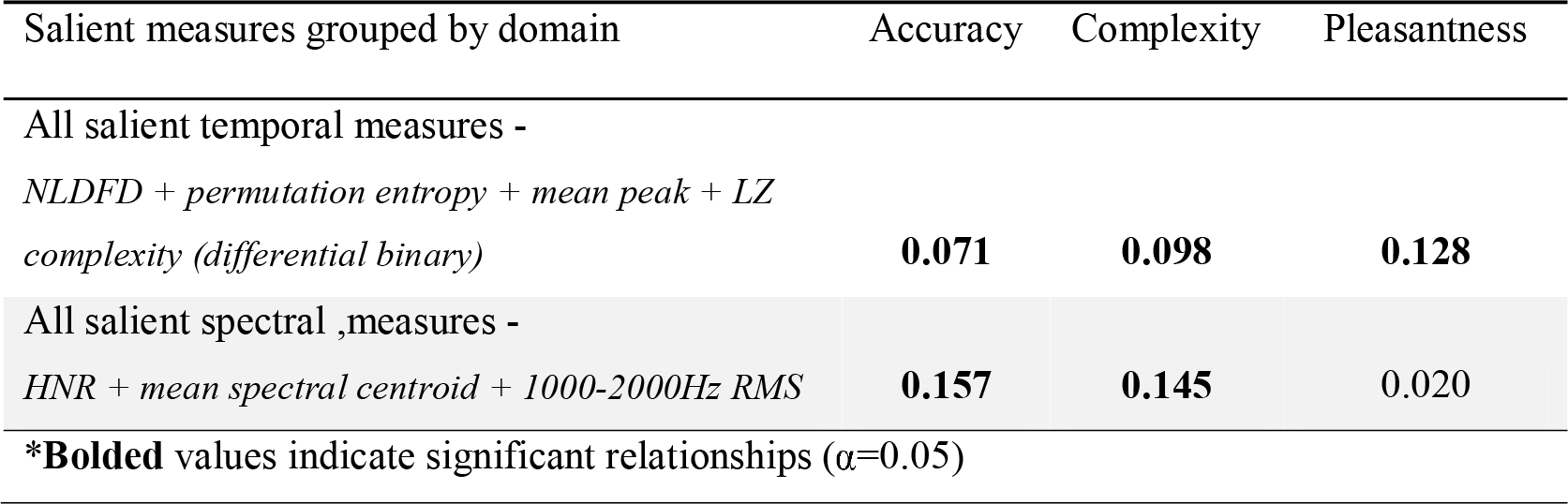
Correlation matrix (R^2^) for Complexity, Pleasantness and identification accuracy, showing their relationships with groups of salient measures.

| Salient measures grouped by domain | Accuracy | Complexity | Pleasantness |
| --- | --- | --- | --- |
| All salient temporal measures -<br>*NLDFD + permutation entropy + mean peak + LZ complexity (differential binary)* | **0.071** | **0.098** | **0.128** |
| All salient spectral measures -<br>*HNR + mean spectral centroid + 1000-2000Hz RMS* | **0.157** | **0.145** | 0.020 |
| * **Bolded** values indicate significant relationships ( $\alpha=0.05$ ) | | | |

The relationships between the measures which best accounted for each of the three perception-related factors were visualised using PCA biplots (Figure 2). With regards to Accuracy of Naming of the NLSs (Figure 2A), HNR and 1000-2000Hz RMS are relatively orthogonal (unaligned) to Accuracy of Naming and the mean spectral centroid lies on the same plane but is almost exactly opposite in direction. With respect to the percept of Complexity of the NLSs, Figure 2B shows that both the 1000-2000Hz RMS measure and HNR are nearly orthogonal to Complexity, and the mean spectral centroid is more closely aligned along the plane of Complexity. Finally, with respect to the percept of Pleasantness of the NLSs, Figure 2C shows that mean peak relative amplitude is the primary measure to align itself roughly with Pleasantness and all of the other salient temporal measures are relatively unrelated, or contribute weakly.

**Figure 2.**
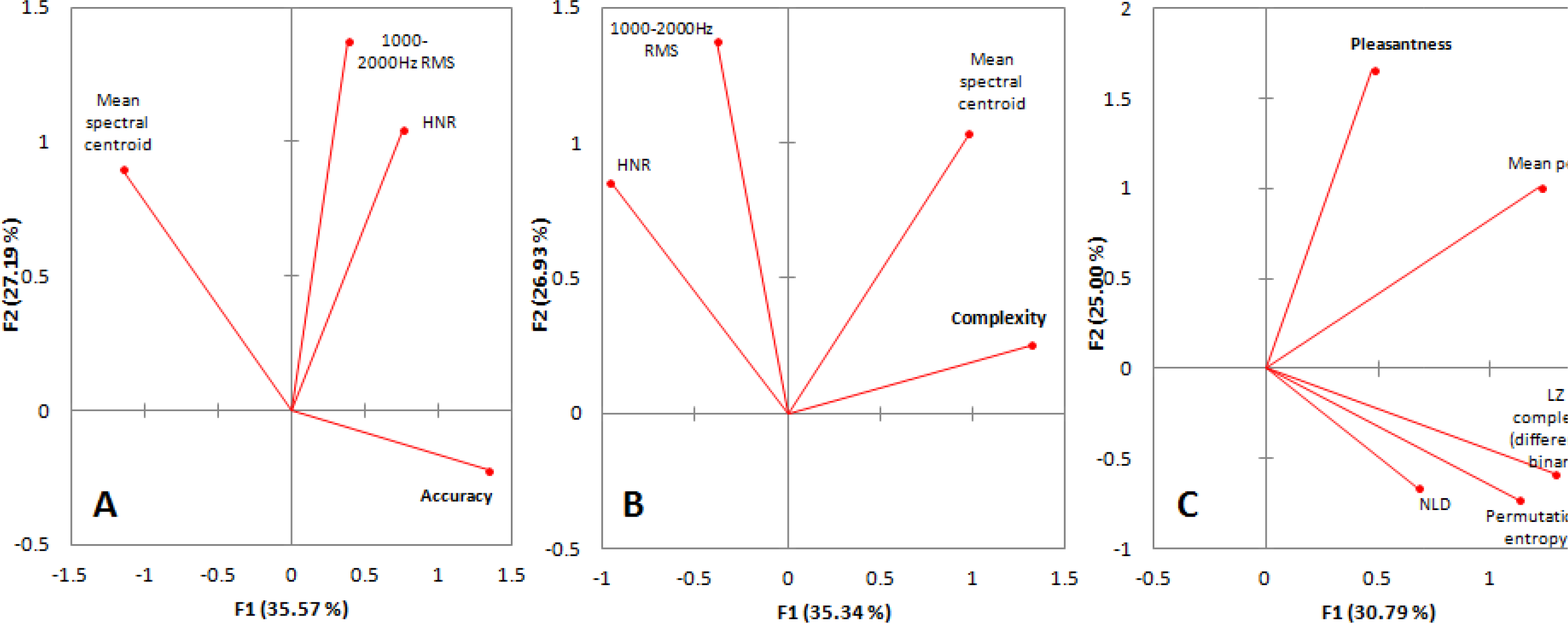
**A:** Biplot of PCA for the salient spectral measures which best described <u>accuracy of naming</u>. MB. Only 62.76% of the variance within these variables is represented. **B:** Biplot of PCA for the salient spectral measures which best described <u>complexity</u>. N.B. Only *82.27%* of the variance within these variables is represented. **C:** Biplot of PCA for the salient spectral measures which best described <u>pleasantness</u>. N.B. Onh 55.79% of the variance within these variables is represented.

#### 3.4.3 Agglomerative hierarchical clustering (AHC)

##### a) AHC of raw values

The resultant clusters of the AHC on the raw *spectral* salient objective measures data for all sounds, were found to have significant (p<0.05) differences in perceptual ratings and Accuracy of Naming a one-way ANOVA and post hoc Tukey’s test, i.e. based upon the objective similarities and differences among all of the NLSs’ in the spectral domain, NLSs formed significantly different groupings in the perceptual domain vis-à-vis differences in Accuracy of Naming and perceptual Complexity. More precisely, these differences were found for Accuracy of Naming (two: p=0.0441; p=0.0441) at the 10-cluster level, for Accuracy of Naming (two: p=0.0014; p=0.0016) and Complexity (two: p=0.0039; p=0.0069) at the five-cluster level, and for Accuracy of Naming (one: p=0.0001) and Complexity (one: p=0.0008) at the three-cluster level. However, the difference at the 10-cluster level for Accuracy of Naming relied on one cluster having only two NLSs. Importantly, no significant differences were found between clusters for Pleasantness at any level of clustering. This is consistent with our previous analyses that spectral measures are strongly related to both Accuracy of Naming and Complexity, but not to Pleasantness (Tables 2-4).

Resultant clusters from the raw *temporal* AHC showed no significant differences at the 10- or four-cluster level. At the two-cluster level there were significant differences for both Accuracy of Naming and Complexity. These differences had, however, much higher p-values (p=0.0306 for Accuracy of Naming and p=0.0470 for Complexity) than those from a comparable cluster level for the spectral AHC, similar for the 3-cluster level (where p=0.0001 for Accuracy of Naming and p=0.0008 for Complexity). This again corroborates with our earlier results showing temporal measures correlate significantly but not as strongly as spectral measures with Accuracy of Naming and Complexity (Tables 2, 3 and 4).

##### b) AHC of principal components

PCAs were conducted using all of the temporal measures and, separately, all of the spectral measures. The first two principal components (PCs), which accounted for the greatest proportions of variance within the original sets of salient measures, were then selected for use in the subsequent (PC) AHC analyses. In the case of the spectral PCA, the first two PCs accounted for 70.78% of the variance, and for the temporal PCA, 61.37% of the original variance was represented in the first two PCs.

Results for the AHC analysis which used the PCs derived from the *spectral* PCA were similar to those from the raw spectral AHC (which used the unmodified data) - there were differences at multiple cluster levels for Accuracy of Naming and Complexity, but not for Pleasantness. The differences were found for Accuracy of Naming (two: p=0.0296; p=0.0167) and Complexity (two: p=0.0218; p=0.0258) at the 10-cluster level, for Accuracy of Naming (two: p=0.0008; p=0.0176) and Complexity (two: p=0.0008; p=0.0176) at the four-cluster level, but for none at the two-cluster level. However, at both the 10- and four-cluster level, the clusters in the ANOVAs for Accuracy of Naming and Complexity has significantly different standard deviations (p<0.05). Thus, these results may be unreliable, however are presented here in the interest of completeness.

Results for the AHC analysis which used the PCs derived from the *temporal* PCA were similar to those from the previous raw temporal AHC - there were differences at multiple cluster levels, including for Accuracy of Naming and Complexity, but at no point were significant differences found between the clusters for Pleasantness. The differences found were for Accuracy of Naming (one: p=0.0491) at the 10-cluster level, for Accuracy of Naming (one: p=0.0296) at the six-cluster level, and for Accuracy of Naming (one: p=0.0027) and Complexity (one: p=0.0115) at the three-cluster level. That significant differences in Pleasantness were not detected in this analysis (as was expected) is possibly due to the fact that only 61.37% of the variance was captured in the temporal PCA. Since, of our salient measures, mean peak relative amplitude is the major or sole contributor to the relationship with Pleasantness and the objective temporal domain, it could be that its contribution was ‘washed-out’ by the lost overall variance or variance contribution from the other salient temporal measures included in the PCA conducted prior to the AHC.

## 4. Discussion

We have made (to our knowledge) the first major attempt at understanding the way humans perceive the Complexity of NLSs in relation to the objective features of those sounds. Two near-orthogonal axes have also been identified in perceptual space and we have added significantly to what we know about the objective determinants of the percepts of Pleasantness, Familiarity, and NLS identification. We also demonstrated the usefulness of AHC analysis on transformed data such as these and how similar methods such as artificial neural networks may help to further tease out complex mappings between objective features of stimuli and their subjective perception as reported by humans.

### 4.1 Relationships among subjective percepts and Accuracy of Naming

As noted earlier, other studies have probed various perceptual properties of NLSs (Halpern et al., 1986; Ballas, 1993; Cycowicz & Friedman, 1998; Marcell et al., 2007; Kumar et al., 2008; Reddy et al., 2009; Reuter & Oehler, 2011; Singh, 2011; Kirmse et al., 2012; Lewis et al., 2012; Talkington et al., 2012) but most focused on only a single percept. Ballas (1993) used the most exhaustive list of perceptual properties, of 22 ratings scales, for 41 NLSs, and condensed this battery of perceptions into three PCs representing 87% of the variance. Given the number of different rating scales, this result shows that subjective ratings can be highly interrelated or interdependent. These ratings did not include Accuracy of Naming, nor did they include Complexity or Pleasantness of NLSs, all of which we considered here. Ballas (1993) did consider Familiarity but the methodology allowed participants to replay the sound as many times as desired and this could affect the other perceptual reports - e.g., Familiarity with an NLS can alter the way it is processed (Cycowicz & Friedman, 1998; Kirmse et al., 2012). Marcell et al. (2007) studied the perceptions of Complexity, Pleasantness, and Familiarity, and the Accuracy of Naming but did not attempt to correlate these perceptual ratings.

We attempted to correlate the percepts of Complexity, Pleasantness, Familiarity, and the Accuracy of Naming. Although all were interrelated to various extents for our dataset of NLSs, there was a dominant relationship between the percepts of Complexity and Familiarity, and Accuracy of Naming: sounds rated as being highly complex are difficult to accurately name and are rated as not familiar, and vice-versa. With the constraint that our subjects were all from a very similar Western industrialised background (albeit of different ethnicities), this indicates that a person’s auditory experience determines their ability to identify NLSs, and ‘complex’ NLSs are rated as such due to a person’s lack of Familiarity with them, independent of any objective characteristics of the sound.

Pleasantness was not well related with the other percepts and especially not with the Accuracy of Naming, except for a weak relationship whereby familiar sounds were rated as slightly more pleasant. Halpern et al. (1986) suggested that the functional purpose of unpleasantness as an auditory feature is to communicate distress or warnings, but this hypothesis was based on similarities between the properties of unpleasant sounds and macaque monkey warning cries, and these similarities are not sufficiently robust to support this hypothesis (Cox, 2008). McDermott & Hauser (2004) found that cotton-top tamarins (*Saguinus oedipus*) had no preference for amplitude-matched white noise versus the sound of a three-pronged metal garden tool scraped down a pane of glass (a ‘screech’ sound comparable to fingernails scraping down a blackboard; Halpern et al., 1986) while humans overwhelmingly preferred the white-noise control (McDermott & Hauser, 2004) despite both species sharing many similarities in perceptual processing for vocalisations (Ramus et al., 2000; Miller et al., 2001a; Miller et al., 2001b; Newport et al., 2004). McDermott & Hauser (2004) also showed that the tamarins significantly preferred looping soundscapes composed of tamarin food chirp sounds versus tamarin distress screams. If the percept of unpleasantness is species-specific, it might not be possible to define this percept in terms of objective properties alone (see below for discussion of the objective properties related to Pleasantness in humans) or may involve semantic or emotive contents which cannot be fully interpreted by another species, or both.

It is interesting to note that highly pleasant sounds, such as nature sounds (Shimai et al. 1993; Marcell et al., 2007; Kumar et al., 2008; Alvarsson et al., 2010), may indicate a relatively safe environment and may impact on health (Cohen et al., 2007). Thus, Alvarsson et al. (2010) show that sounds such as nature sounds (consistently rated as highly pleasant; Shimai et al. 1993; Marcell et al., 2007; Kumar et al., 2008; Alvarsson et al., 2010), facilitate recovery from sympathetic activation in humans after experiencing a psychological stressor but that other sounds, such as road traffic sounds (often rated as unpleasant; Marcell et al., 2007; Alvarsson et al., 2010), do not facilitate the same recovery. However, we recognise that this facilitation or absence of recovery may also involve higher-order emotional or subjective elements and not just be due to the objective features of these NLSs.

### 4.2 Relationships among objective measures of NLSs

Many correlations were found among the objective measures in both temporal and spectral domains, some expected and some not. We acknowledge that these relationships may only be true for our dataset of NLSs and may not be a general rule for all types of NLSs, let alone all types of signals.

Although dissimilar in methodology, both fractal dimension estimation techniques - Higuchi and NLD - were highly correlated, as also seen when these analyses were originally applied to EEG waveforms (Kalauzi et al., 2009). Interestingly, we have reported the first significant correlation between a fractal dimension estimate and the LZ complexity measure (average binary and modified zone binary). Since both measures ultimately attempt to find self-similarity or repeating aspects in a signal, it is not unexpected that they carry some common information. However, the degree of common information was not always similar, e.g., the Higuchi FD correlated poorly with LZ complexity (modified zone binary) (r=0.361; Table 2) whereas the NLD FD was more correlated (r=0.577; Table 2).

The most significant relationship among the spectral measures, the negative relationship between the SFM and HNR, is logical since if a sound is highly harmonic it cannot also be ‘flat’ in its spectrum. The fact that the SFM and SSI are related explains why they were also well correlated and therefore why there was also a relationship between SSI and HNR.

### 4.3 Categories of sounds differentiated by source were not strongly differentiated by either subjective percepts or objective measures

We analysed our NLSs database for homogeneity to examine if stronger relationships could be found by removing outlier NLSs which did not fit a given trend between an objective measure and subjective percept or Accuracy of Naming. We sought to determine if different categories of NLSs followed their own trends, independently of other categories (see 2.4.2).

With respect to Accuracy of Naming, non-linguistic human vocalisations (which we classed under ‘primates’) were the most accurately identified NLS category, although only the ‘other’ category was significantly less so (p=0.0037 c.f. ‘primates’). This differs slightly from the results of Inverso & Limb (2010), who found ‘mechanical/alerting’ sounds to be slightly more recognisable than ‘human’ sounds. However, their ‘human’ category also included non-vocalisations like the sound ‘footsteps’ and their study participants were experienced cochlear implant (CI) users, not normal-hearing university students. It is not known if the perception of NLSs is the same between the two categories of people, but these findings raise caveats about assuming that the understanding of the perception of NLSs by normal-hearing subjects can directly translate to the perception of NLSs by deaf subjects using a CI.

With respect to perceived complexity, primate sounds in our database were rated as significantly less complex than the sounds of other animals; this may reflect the similarity between human sounds and primate sounds (Hauser & Fitch, 2003), and therefore be a Familiarity factor.

For ratings of Pleasantness, for our NLS categories, nature and music sounds were rated as the most pleasant, as in previous studies (Shimai et al. 1993; Marcell et al., 2007; Kumar et al., 2008; Alvarsson et al., 2010).

Finally, the only salient objective measures showing differences among the different NLS source categories were LZ complexity (differential binary) and the HNR. Previous studies (Gygi et al., 2007; Leaver & Rauschecker, 2010; Lewis et al. 2012) have noted the importance of harmonicity to NLS classification but did not determine if there were differences among different experimenter-determined categories of sounds for HNR (or LZ complexity).

Overall, there were some significant differences among different categories of NLS sources for subjective percepts and for objective measures of sounds but there were many groups for which no significant differences were observed for either. Indeed, Figures 5 and 6 show that objective features of sounds can be similar or even identical between different source categories. Later categorisation of the different sounds using AHC analyses showed that this method of objective categorising was more revealing of similarities and differences between different sounds (note: not different sound categories) for Complexity and the Accuracy of Naming. Thus, identifying a sound’s source appears only partly to rely on objective measures, and other information, perhaps visual integrative learning or memory (Thompson & Allan, 1994; Giard & Peronnet, 1999), may also play a role. Higher-order associations may also provide context and input into perceptions of Complexity and Pleasantness, as for other perceptions (Johnson et al. 1999).

**Figure 4.**
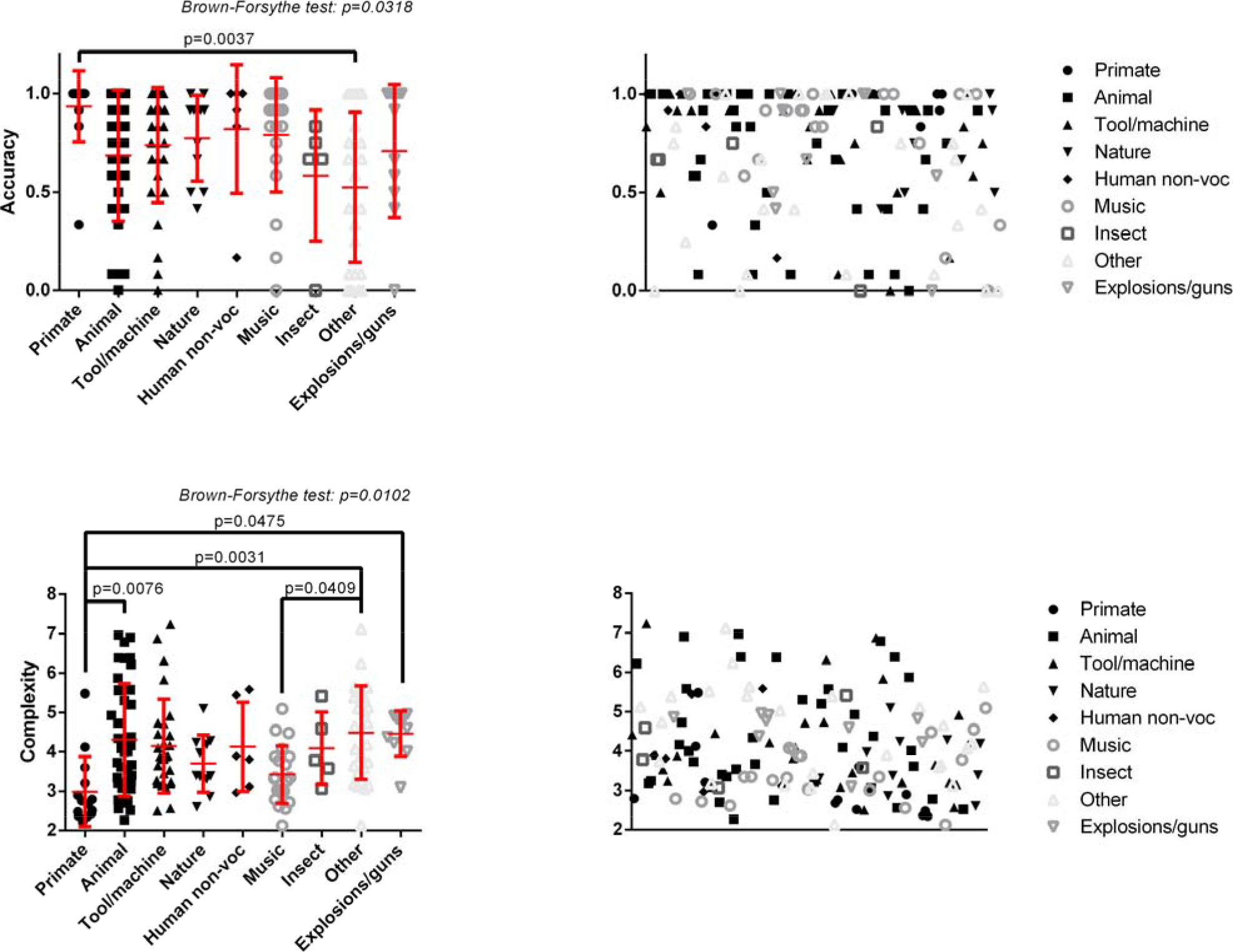
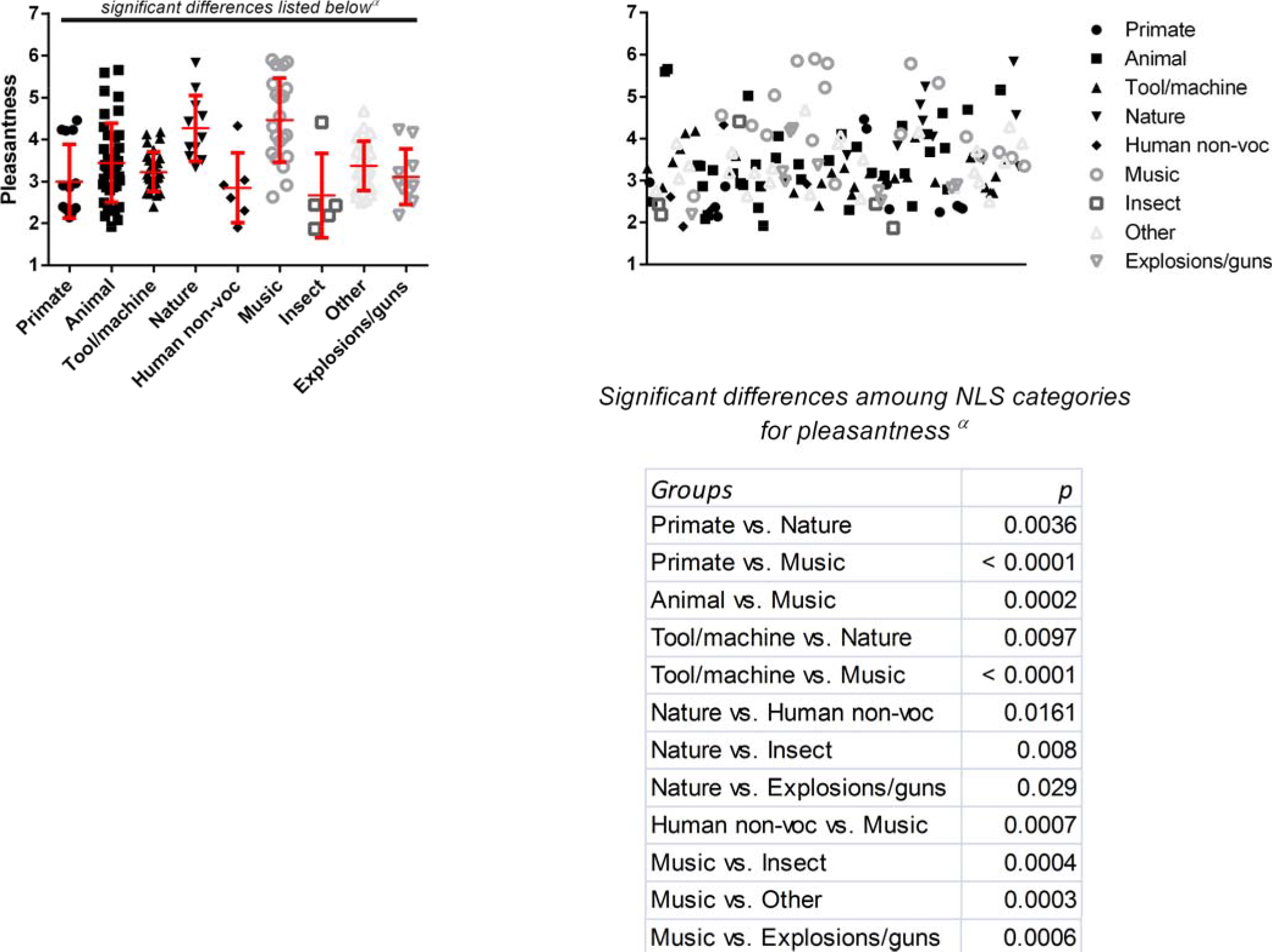
Spreads for the percepts and accuracy are shown here for each of the nine experimenter-determined NLS categories based on sound source. Significant differences between categories were found in each of accuracy, complexity and pleasantness, with the greatest differences found in pleasantness. The ANOVAs for both accuracy and complexity had significantly different standard deviations among their groups, whereas the ANOVA for pleasantness did not.

**Figure 5.**
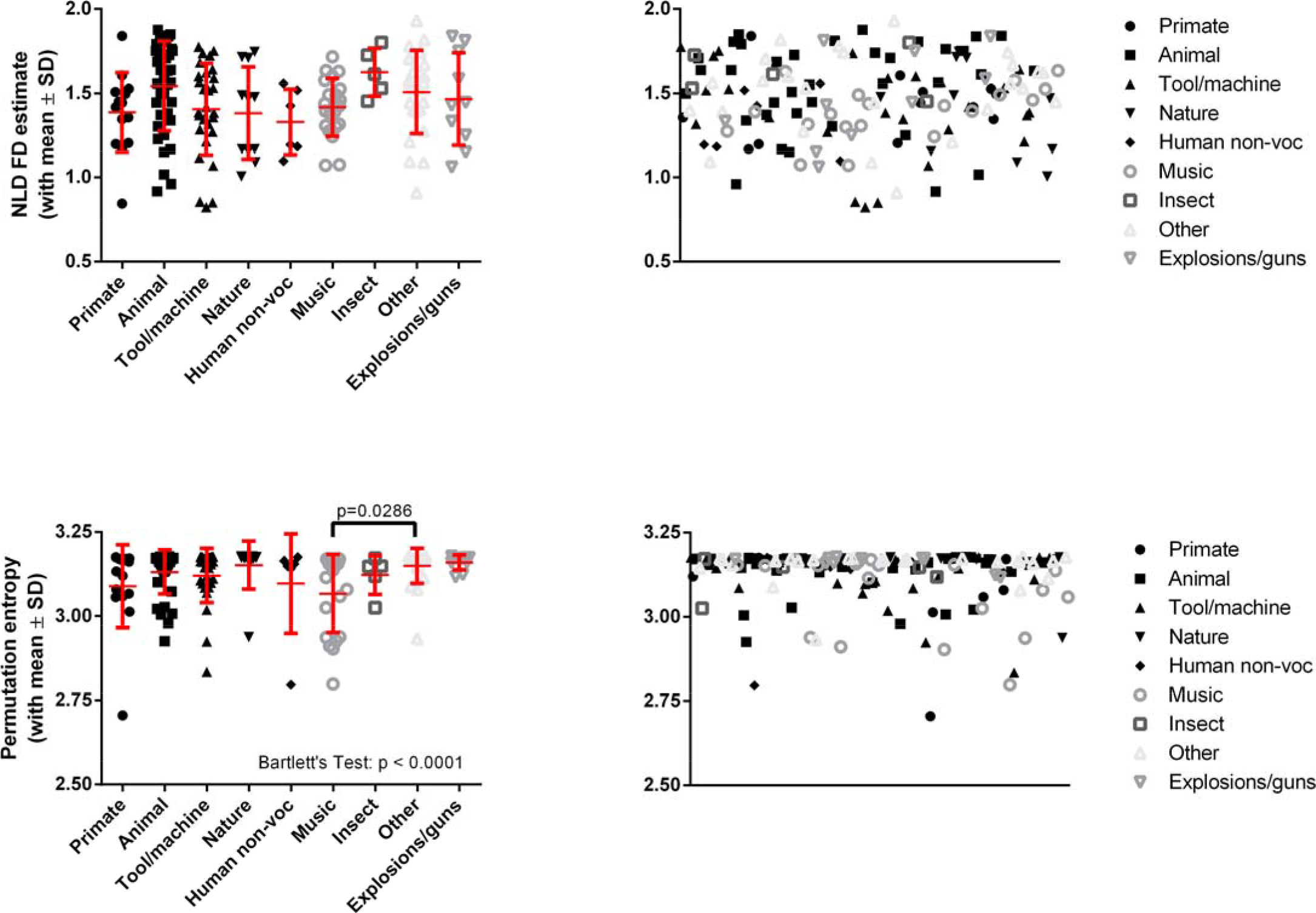
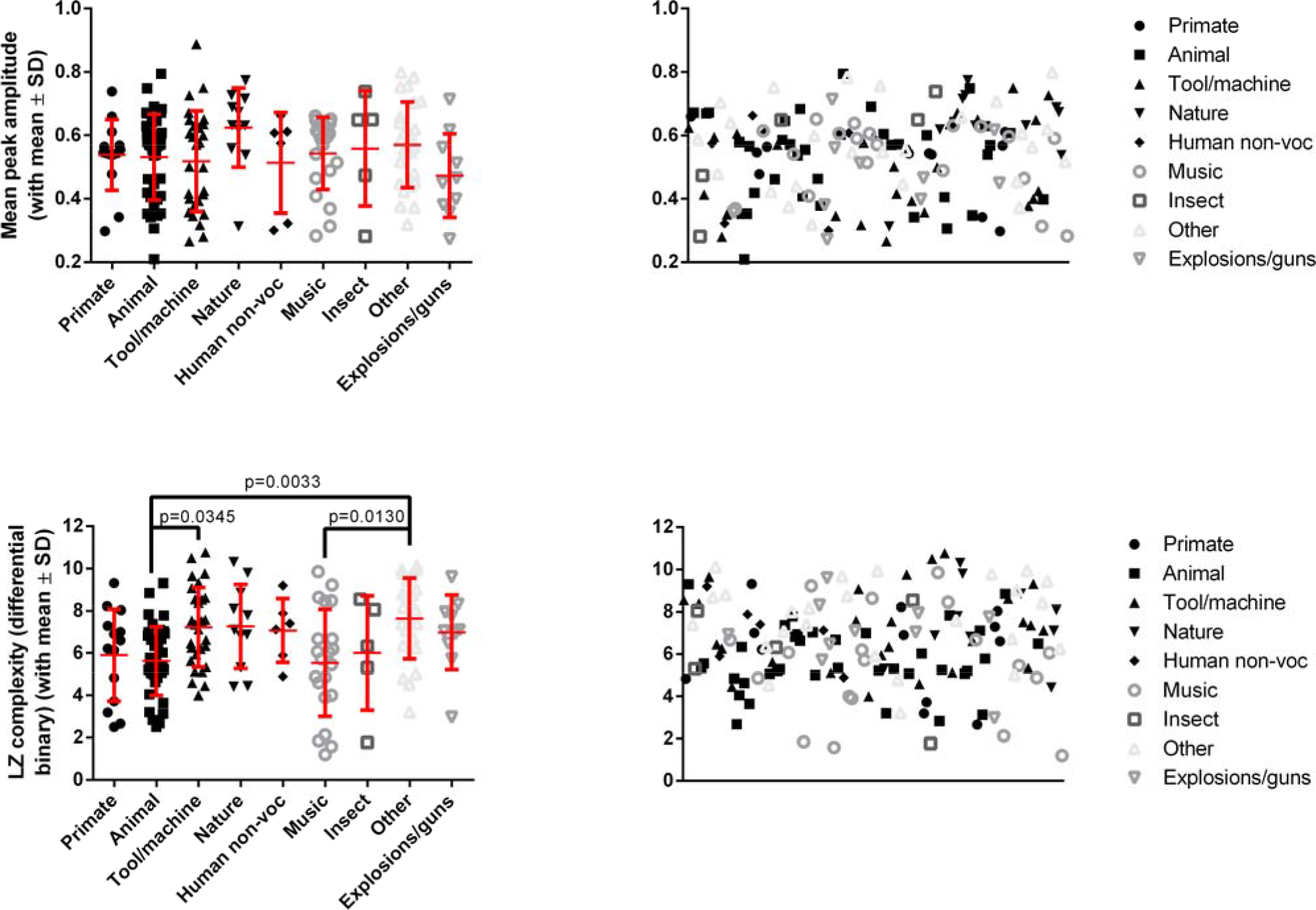
Spreads for each of the temporal salient measures are shown here for each of the nine experimenter-determined NLS categories based on sound source. Significant differences between categories were found in the cases of permutation entryopy and the LZ complexity measure using differential binary. However, the only common dissimilarity of these two measures was between the ‘Music’ and ‘Other’ NLS categories.

**Figure 6.**
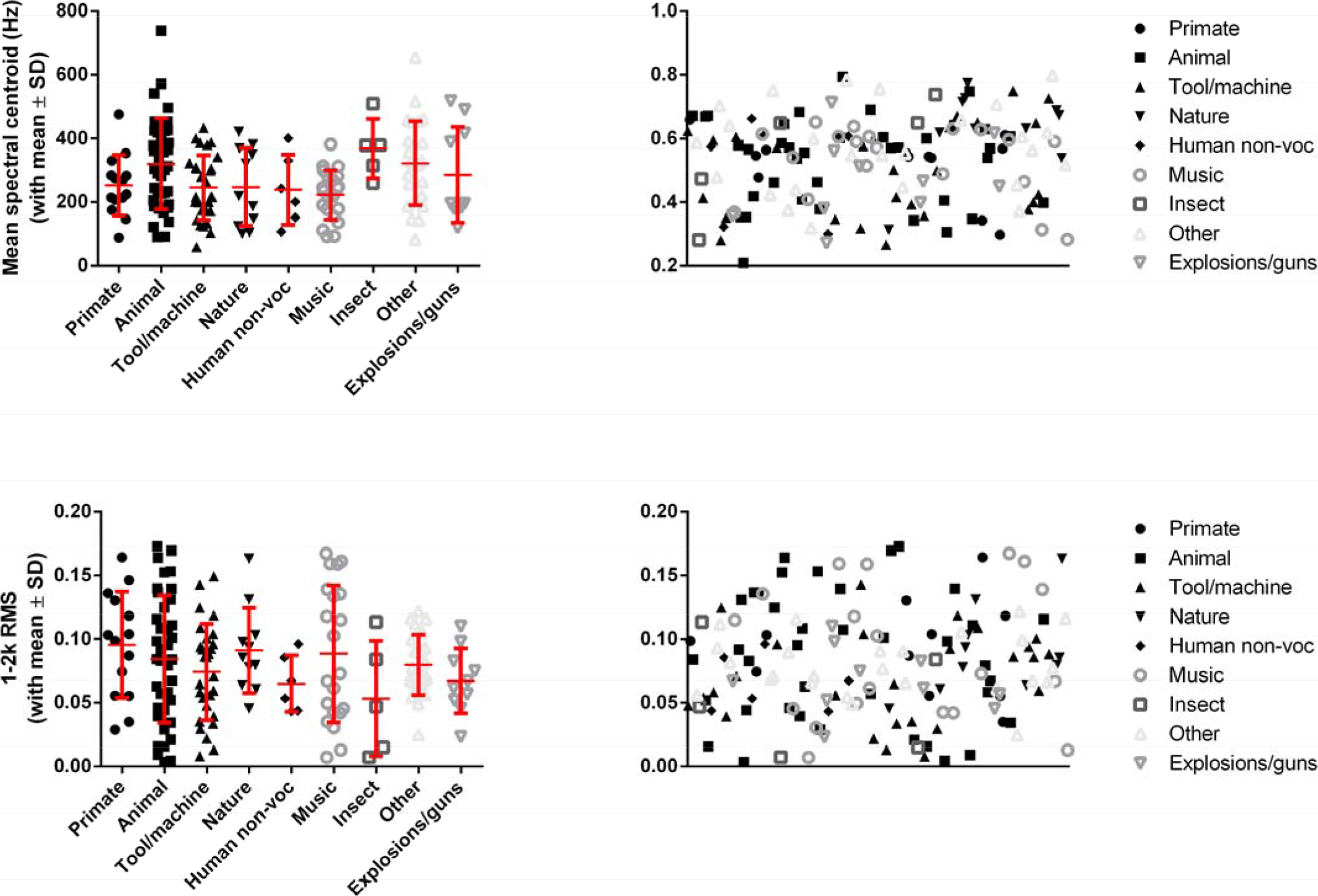
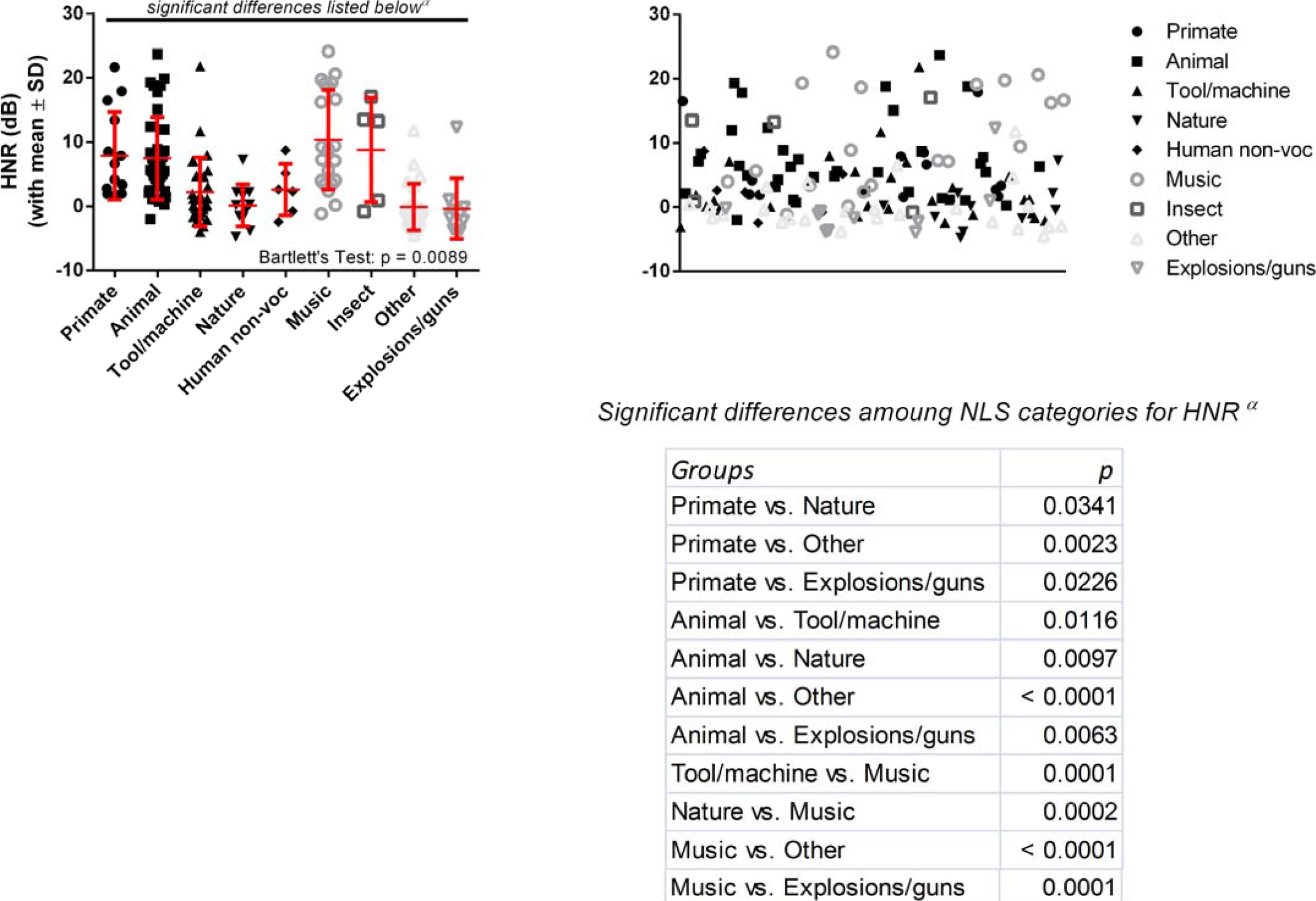
Spreads for each of the spectral salient measures are shown here for each of the nine experimenter-determined NLS categories based on sound source. Significant differences between categories were found only in the HNR values. Compared to the temporal salient measures which showed some significant differences amoung the categories, there were many more differences for HNR. This highlighted HNR’s importance moving forward.

This conclusion has implications for the identification of NLSs by CI users (viz., Inverso & Limb, 2010). The poor success of CI users in identifying NLSs may have less to do with the objective properties of the NLSs and more to do with other attributes, e.g., remembering and associating that sound with an image, based on experience or emotional valency. Work on the perceptual learning of spectrally degraded speech and NLSs (Loebach & Pisoni, 2008) shows that speech training does not generalise to NLSs but that NLS training *does* generalise to speech. This suggests that the development of specific NLS training programs for CI or hearing aid users would be greatly beneficial, both to allow recognition of an important set of everyday sounds with strong emotional or survival value and to feed into speech recognition.

### 4.4 Complexity, Familiarity and Accuracy of Naming

Since different categories of sounds, segregated by sound source, were not strongly differentiated by either subjective percepts or objective measures, we used the entire NLSs database to find correlations with the objective measures along our two, near-orthogonal perceptual dimensions of Complexity and Pleasantness. Since Familiarity closely matched Complexity and the Accuracy of Naming, any relationship which was true for one would often exist oppositely for the other. (Pleasantness was a near-orthogonal perceptual dimension to the Complexity-Familiarity-Accuracy-of-Naming dimension, and its relationships with objective measures are discussed in 4.5.)

The consistent correlation between objective measures and Familiarity suggests either (1) our subjects controlled their auditory experience with respect to objective features of NLSs (e.g., they avoided sounds with high FDs, and thus were unfamiliar with them) or (2) there is something intrinsic to these objective features which makes NLSs easier to become familiar with. The latter hypothesis seems more plausible.

Our standard multivariate analyses showed that both temporal and spectral measures can be associated significantly with differences in Complexity and the Accuracy of Naming (Tables 5 and 6). When the sounds in our database were separated based on these objective measures using AHC analyses (and incorporating PCA), differences arose between the resultant clusters for both Complexity and the Accuracy of Naming. However, the resultant clusters from AHC analyses which included only spectral measures tended to show greater numbers of and more significant differences in Complexity and Accuracy of Naming. This suggests humans may assess spectral features, more than temporal features, when assessing NLS Complexity, Familiarity, or when seeking identify it. Favouring spectral features over temporal ones may be a more efficient means of neural processing insofar as being able to more readily make a subjective judgement about a NLS, since relying on temporal features may require a longer time exposure to the NLS.

### 4.5 Pleasantness

For ratings of Pleasantness, temporal measures or combinations thereof had the highest explanatory power and none of the salient spectral measures were significantly correlated - alone or in combination - with Pleasantness; clusters created using spectral measures did not show any significant differences in Pleasantness. The fact that temporal information becomes highly important in CI users with spectrally degraded stimuli (Fu et al., 2004), and because a decreased level of music appreciation is found in the same individuals (Nimmons et al., 2008; Philips et al., 2012), makes it likely that temporal features are important to the appreciation of sounds as being pleasant. Poorer temporal resolution might mean that such features are indiscernible to a person with hearing loss (Francart et al., 2015), and thus result in a lowered appreciation of music.

However, past studies have repeatedly shown the importance of spectral features to percepts of *Unpleasantness* in NLSs (Halpern et al., 1986; Cox, 2008; Kumar et al., 2008; Reuter & Oehler, 2011). This might be because *Unpleasantness* has different objective drivers than does *Pleasantness,* i.e., the presence of an objective feature might make something unpleasant but its absence might have a neutral or null effect on Pleasantness. However, it must also be recognised that there are significant methodological differences between our study and the previous studies: Halpern et al. (1986) and (Rueter & Oehler, 2011) used 16 digitally resynthesised and filtered NLSs; Kumar et al. (2008) used 75 auditory representations of NLSs in a modelled primary auditory cortex (Shamma, 2003); and Cox (2008) used 34 NLSs but did not measure any objective features. Further, none of these studies tested the main temporal measure which we found to correlate with Pleasantness - mean peak relative amplitude. Based on the latter difference we would argue that these studies do not negate our finding that Pleasantness perception is based on temporal measures. This is not to say that it has no spectral bases, but rather that it at least has some temporal ones.

### 4.6 Conclusions

This study represents a first and critical step to the principled subjective and objective characterisation of NLSs, and could allow NLSs libraries to be used in the evaluation or training of people with hearing or cognitive impairments in the processing of NLSs or their features. We have shown that a variety of objective measures and percepts (including the Accuracy of Naming) have strong interrelationships for our NLSs database, and we have identified two near-orthogonal dimensions in perceptual space. One practical implication of this database is in the creation of a NLSs hearing test or training regime. Information found by a NLSs hearing test could identify if a person finds temporal or spectral information more difficult to process, and this (married with a NLS training regiment) could aid clinicians in developing more specific treatments and rehabilitation strategies for their patients. Another potential benefit could be in the identification and removal of NLSs where they represent unwanted background noise, such as in imperfect binary algorithms to digitally separate speech from noise. Such algorithms could be incorporated into hearing aids or CIs to dramatically counteract one the most commonly complained symptoms of hearing loss (the cocktail party effect).

## 6. Appendices / Supplementary Material

### Appendix I: List of sounds and assigned sound type categories

|  | *Percentage*<br>*of data set* | *Number of sounds* |
| --- | --- | --- |
| **Primate (#1)** | 8.86 | 14 |
| **Animal (#2)** | 24.05 | 38 |
| **Tool/machine (#3)** | 17.72 | 28 |
| **Nature (#4)** | 6.96 | 11 |
| **Human non-voc (#5)** | 3.80 | 6 |
| **Music (#6)** | 13.29 | 21 |
| **Insect (#7)** | 3.16 | 5 |
| **Other (#8)** | 15.82 | 25 |
| **Explosions/guns (#9)** | 6.33 | 10 |
| *Total* |  | *158* |

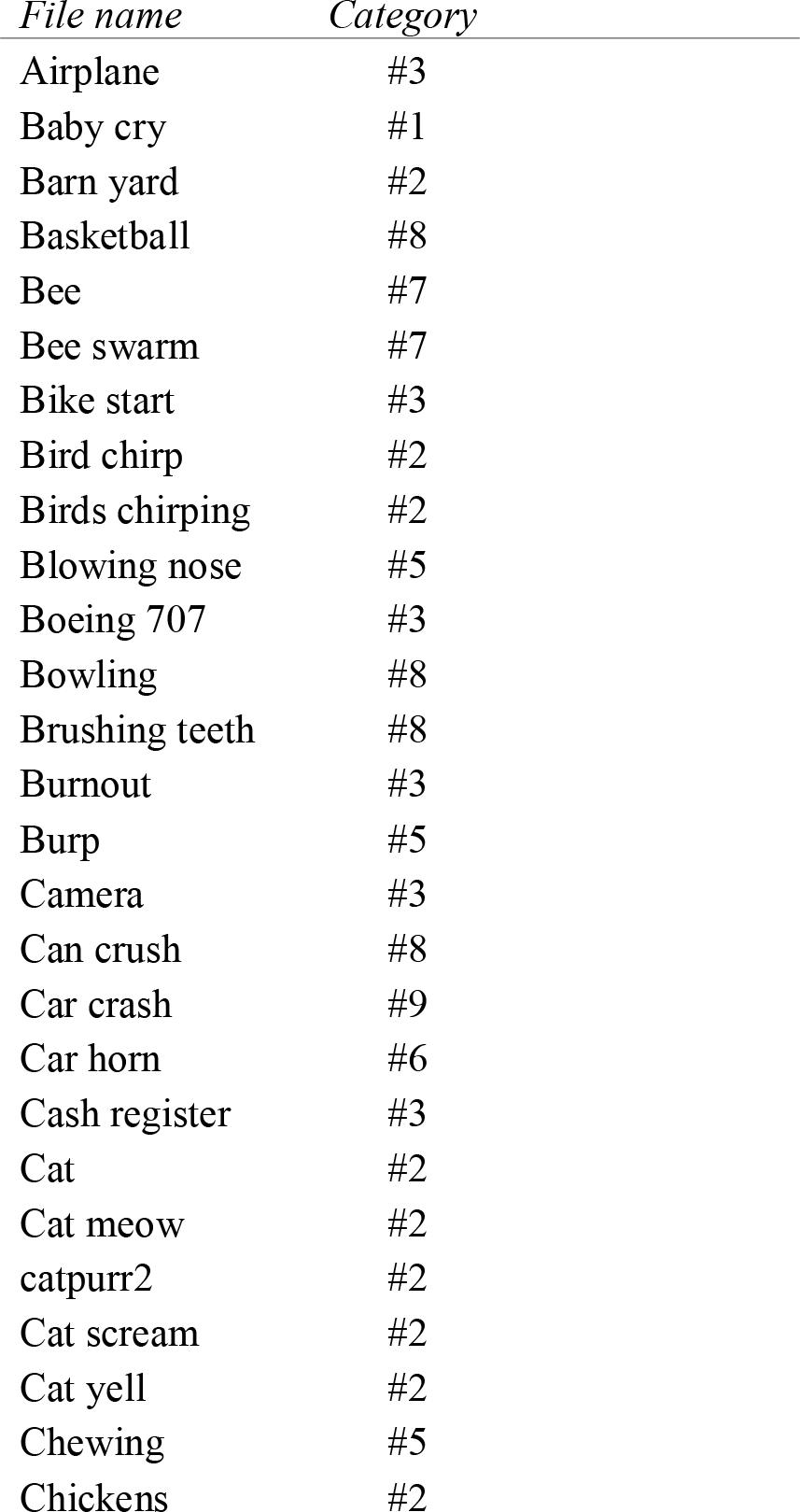

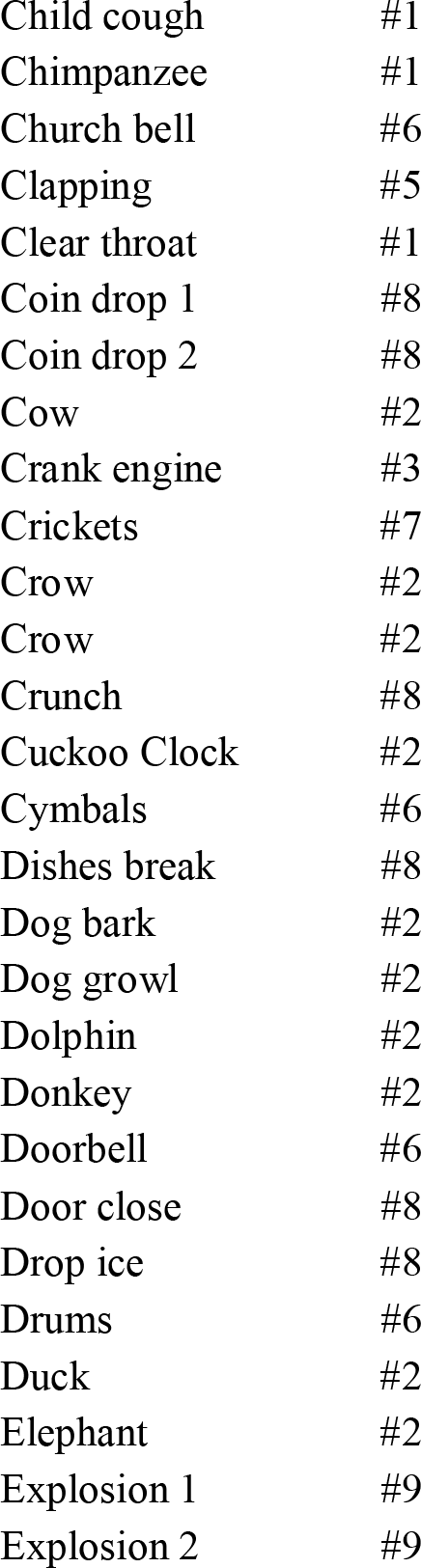

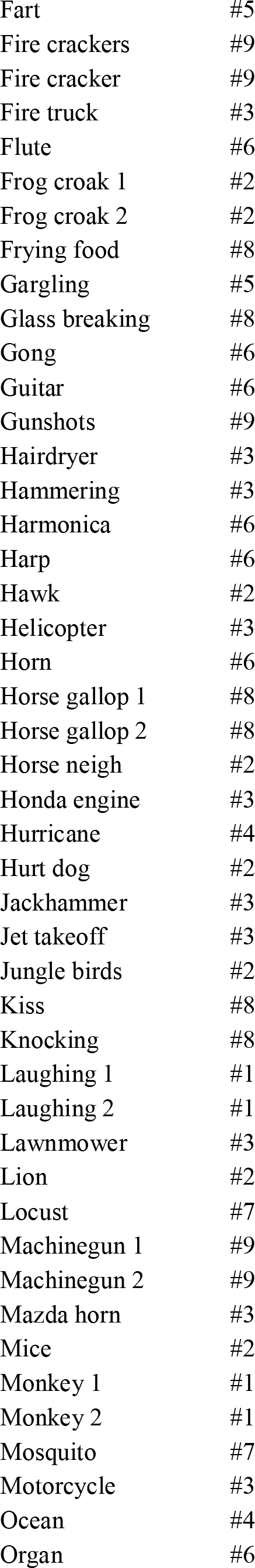

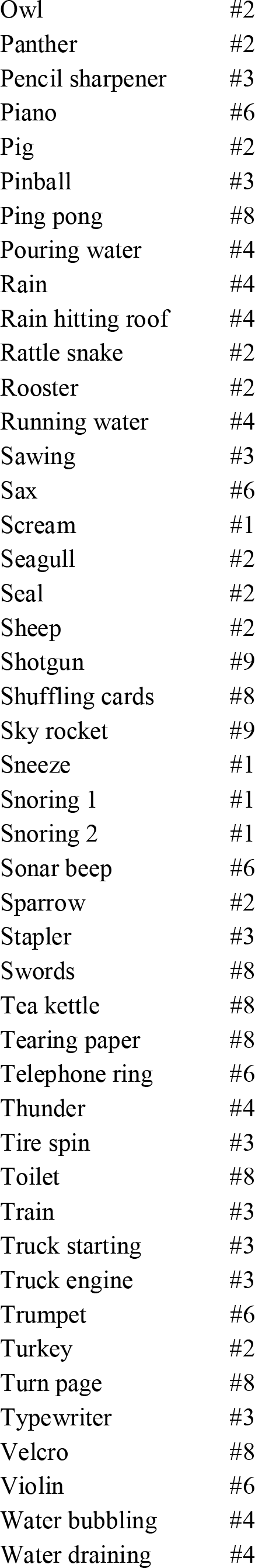

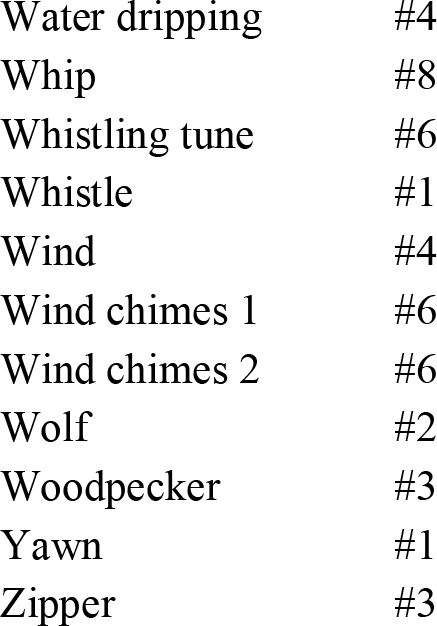

### Appendix II Identification and selection of salient objective measures for NLSs

As expected, many measures which were theoretically related or which measured similar qualities of an NLS were significantly correlated with one another, e.g. HNR and SFM (r=-0.822); Higuchi and NLD estimates (r=0.812). In these instances choosing just one of these inter-related measures would provide nearly the same amount of information about a NLS as choosing both for the subsequent analyses. Although there are many significant relationships among the measures, some of them are quite weak, e.g. Higuchi FD estimate and duration are significantly but weakly correlated (r=0.059). Therefore, the choice of measures should include some consideration towards the strength of these relationships. However, any such consideration would have to be made on an entirely arbitrary basis since the ‘strength’ of a correlation is subjective and relative.

The arbitrary cut-off point chosen was r=±0.45, i.e., if the correlation between two measures was ≥±0.45, only one of the pair would be included in subsequent analyses. Ten cross-relationships with r≥±0.45 were found among the temporal measures and seven such cross-relationships among the spectral measures. Retaining only one from each such cross-relationship reduced our original set of 19 measures to a subset of seven.

**App II. Table 1.** Correlation matrix (Pearson) for all temporal measures, showing some significant levels of covariance between some measures.

| Variables | HFD | NLDFD | Duration | Peaks /<br>sec | Mean<br>peak | SD of<br>peaks | Sample<br>entropy | Permutation<br>entropy | LZ<br>complexity<br>(average<br>binary) | LZ<br>complexity<br>(modified<br>zone binary) | LZ<br>complexity<br>(differential<br>binary) |
| --- | --- | --- | --- | --- | --- | --- | --- | --- | --- | --- | --- |
| HFD |  | 0.812 | 0.059 | 0.143 | 0.202 | 0.256 | -0.193 | 0.115 | 0.66 | 0.361 | 0.277 |
| NLDFD | **0.812** |  | 0.009 | 0.233 | 0.085 | 0.165 | 0.019 | 0.112 | 0.697 | 0.577 | 0.126 |
| Duration | 0.059 | 0.009 |  | -0.242 | 0.155 | 0.206 | -0.008 | -0.04 | 0.164 | 0.055 | 0.18 |
| Peaks/sec | 0.143 | 0.233 | -0.242 |  | -0.568 | -0.518 | 0.261 | -0.116 | -0.024 | 0.049 | -0.149 |
| Mean peak | 0.202 | 0.085 | 0.155 | **-0.568** |  | 0.9 | -0.691 | 0.127 | 0.334 | -0.074 | 0.237 |
| SD of peaks | 0.256 | 0.165 | 0.206 | **-0.518** | **0.9** |  | -0.63 | 0.204 | 0.419 | 0.012 | 0.237 |
| Sample entropy | -0.193 | 0.019 | -0.008 | 0.261 | **-0.691** | -0.63 |  | -0.019 | -0.225 | 0.403 | -0.242 |
| Permutation<br>entropy | 0.115 | 0.112 | -0.04 | -0.116 | 0.127 | 0.204 | -0.019 |  | 0.419 | 0.349 | 0.305 |
| LZ complexity<br>(average binary) | **0.66** | **0.697** | 0.164 | -0.024 | 0.334 | 0.419 | -0.225 | 0.419 |  | 0.614 | 0.604 |
| LZ complexity<br>(modified zone<br>binary) | 0.361 | **0.577** | 0.055 | 0.049 | -0.074 | 0.012 | 0.403 | 0.349 | **0.614** |  | 0.363 |
| LZ complexity<br>(differential<br>binary) | 0.277 | 0.126 | 0.18 | -0.149 | 0.237 | 0.237 | -0.242 | 0.305 | **0.604** | 0.363 |  |

**App II. Table 2.**
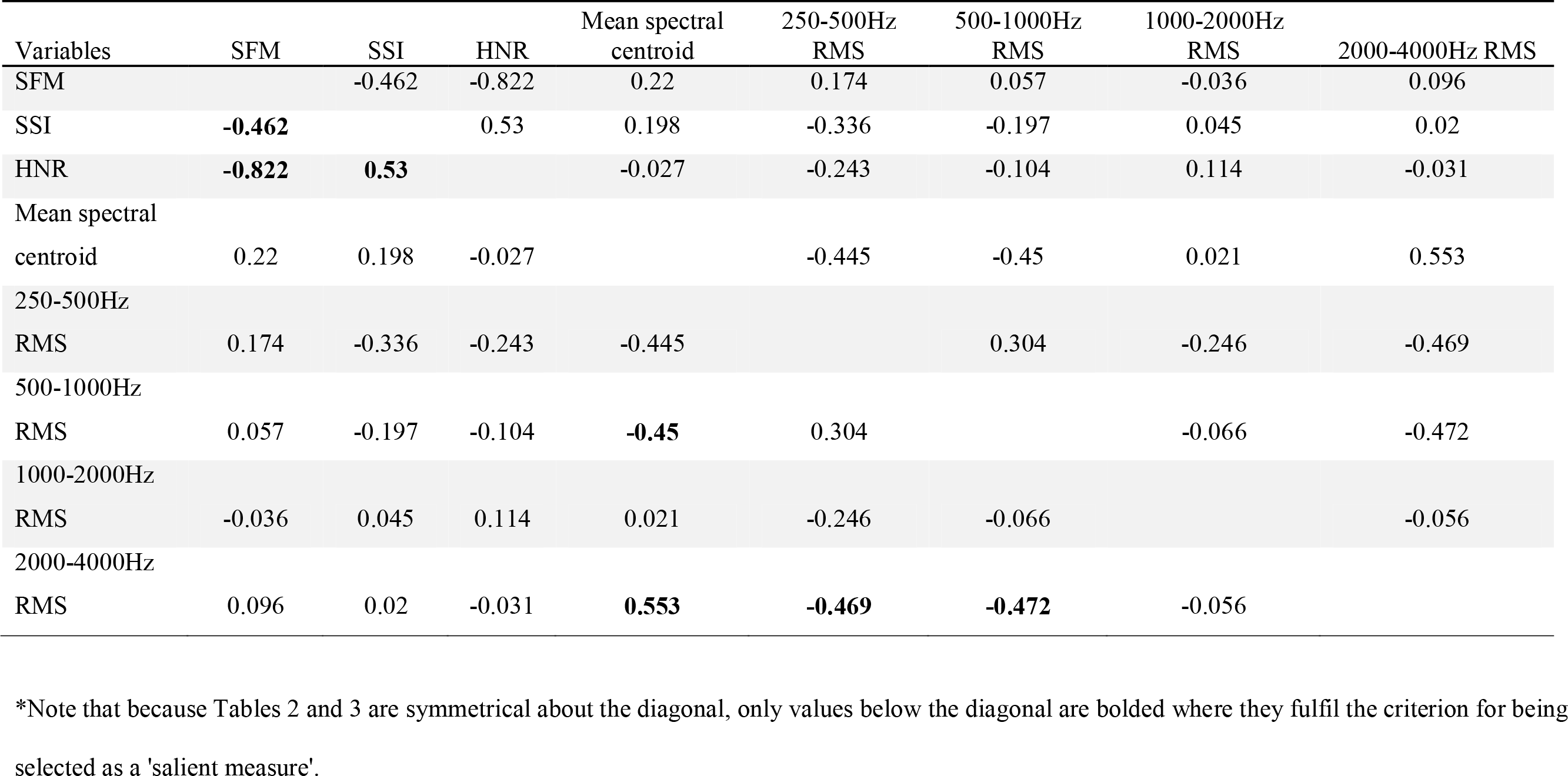
Correlation matrix (Pearson) for all spectral measures, showing some significant levels of covariance between some measures.

| Variables | SFM | SSI | HNR | Mean spectral<br>centroid | 250-500Hz<br>RMS | 500-1000Hz<br>RMS | 1000-2000Hz<br>RMS | 2000-4000Hz RMS |
| --- | --- | --- | --- | --- | --- | --- | --- | --- |
| SFM |  | -0.462 | -0.822 | 0.22 | 0.174 | 0.057 | -0.036 | 0.096 |
| SSI | **-0.462** |  | 0.53 | 0.198 | -0.336 | -0.197 | 0.045 | 0.02 |
| HNR | **-0.822** | **0.53** |  | -0.027 | -0.243 | -0.104 | 0.114 | -0.031 |
| Mean spectral<br>centroid | 0.22 | 0.198 | -0.027 |  | -0.445 | -0.45 | 0.021 | 0.553 |
| 250-500Hz<br>RMS | 0.174 | -0.336 | -0.243 | -0.445 |  | 0.304 | -0.246 | -0.469 |
| 500-1000Hz<br>RMS | 0.057 | -0.197 | -0.104 | **-0.45** | 0.304 |  | -0.066 | -0.472 |
| 1000-2000Hz<br>RMS | -0.036 | 0.045 | 0.114 | 0.021 | -0.246 | -0.066 |  | -0.056 |
| 2000-4000Hz<br>RMS | 0.096 | 0.02 | -0.031 | **0.553** | **-0.469** | **-0.472** | -0.056 |  |

\*Note that because Tables 2 and 3 are symmetrical about the diagonal, only values below the diagonal are bolded where they fulfil the criterion for being selected as a 'salient measure'.

### Appendix III NLSs database heterogeneity

